# EPAC2 is required for corticotropin-releasing hormone-mediated spine loss

**DOI:** 10.1101/607598

**Authors:** Zhong Xie, Peter Penzes, Deepak P. Srivastava

**Affiliations:** Department of Physiology, Feinberg School of Medicine, Northwestern University, Chicago, IL; Department of Psychiatry and Behavioral Sciences, Feinberg School of Medicine, Northwestern University, Chicago, IL; Center for Autism and Neurodevelopment, Northwestern University, Chicago, IL; Department of Basic and Clinical Neuroscience, Maurice Wohl Clinical Neuroscience Institute, Institute of Psychiatry, Psychology and Neuroscience, King’s College London, London, SE5 9RT, UK; MRC Centre for Neurodevelopmental Disorders, King’s College London, London SE1 1UL, UK

**Author notes:** Correspondence: Peter Penzes, Department of Physiology, Feinberg School of Medicine, Northwestern University, Chicago, IL, USA,; or Deepak Srivastava, Department of Basic and Clinical Neuroscience, King’s College London, London, SE5 9RT, UK.

**Keywords:** Stress, dendritic spine, excitatory synapse, cortical neurons, Rap, small GTPase

## Abstract

Corticotropin-releasing hormone (CRH) is produced in response to stress. This hormone plays a key role in mediating neuroendocrine, behavioral, and autonomic responses to stress. The CRH receptor 1 (CRHR1) is expressed in multiple brain regions including the cortex and hippocampus. Previous studies have shown that activation of CRHR1 by CRH results in the rapid loss of dendritic spines. Exchange protein directly activated by cAMP (EPAC2, also known as RapGEF4), a guanine nucleotide exchange factor (GEF) for the small GTPase Rap, has been linked with CRHR1 signaling. EPAC2 plays a critical role in regulating dendritic spine morphology and number in response to several extracellular signals. But whether EPAC2 links CRHR1 with dendritic spine remodeling is unknown. Here we show that CRHR1 is highly enriched in the dendritic spines of primary cortical neurons. Furthermore, we find that EPAC2 and CRHR1 co-localize in cortical neurons. Critically, short hairpin RNA-mediated knockdown of *Epac2* abolished CRH-mediated spine loss in primary cortical neurons. Taken together, our data indicate that EPAC2 is required for the rapid loss of dendritic spines induced by CRH. These findings identify a novel pathway by which acute exposure to CRH may regulate synaptic structure and ultimately responses to acute stress.

## Introduction

Corticotropin-releasing hormone (CRH) is a 41-amino acid neuropeptide that is an important regulator of hormonal, behavioral, and autonomic responses to stress (Henckens *et* al., 2016; Sanders & Nemeroff, 2016). CRH is expressed in discrete regions within the central nervous system (Grammatopoulos, 2012; Henckens *et al.*, 2016), and CRH receptors are expressed in multiple brain regions (Henckens *et al.*, 2016). The CRH receptor type 1 (CRHR1) is a 7-transmembrane G-protein coupled receptor that transmits signals via the Gsα-mediated regulation of cAMP (Traver *et al.*, 2006; Grammatopoulos, 2012). The CRH peptide has been shown to be released locally within the amygdala, hippocampus, and cortex, and to be involved in the modulation of cognition, including memory and anxiety, during stress (Henckens *et al.*, 2016; Sanders & Nemeroff, 2016). Moreover, chronic exposure to CRH may have long-lasting detrimental effects (Maras & Baram, 2012; Henckens *et al.*, 2016). Interestingly, acute exposure to CRH also results in the loss of dendritic spine on CA1 hippocampal neurons; this effect could be blocked by CRHR1 antagonism (Chen *et al.*, 2008; Andres *et al.*, 2013). CRH-mediated spine loss in the hippocampus have been linked with RhoA signaling (Chen *et al.*, 2013), as well as a nectin-3/afadin complex (Wang *et al.*, 2013).

Exchange protein directly activated by cAMP 2 (EPAC2, also known as cAMP-GEFII or RapGEF4) is a signaling protein present in forebrain postsynaptic densities (Woolfrey *et al.*, 2009; Srivastava *et al.*, 2012). This protein has been shown to be involved in a range of cognitive function including social behaviors (Srivastava *et al.*, 2012) and learning and memory (Yang *et al.*, 2012). Interestingly, recent studies have also implicated the *EPAC2* gene in the response to stress, anxiety, and depression (Zhou *et al.*, 2016; Aesoy *et al.*, 2018). Interestingly, EPAC2 has been suggested to mediate CRH/CRHR1 coupling to the ERK-MAPK pathway (Traver *et al.*, 2006; Grammatopoulos, 2012; Inda *et al.*, 2016). Moreover, EPAC2 has also been shown to regulate dendritic spine morphology, motility, and density (Woolfrey *et al.*, 2009; Srivastava *et al.*, 2012). Based on these previous studies, we hypothesized that EPAC2 might play a role in CRH/CRHR1-mediated spine alterations.

Here we investigated the presence of CRHR1 at dendritic spines of primary cortical neurons. We examined whether CRHR1 and EPAC2 co-localized within cortical neurons and whether acute exposure to CRH altered spine density in cortical neurons. Finally, we tested whether EPAC2 was required for CRH-mediated spine loss.

## Materials and Methods

### Reagents

CRH was purchased from Bio-Techne (“CRF”, Cat. No. 1151). Antibodies used: green fluorescent protein (GFP) mouse monoclonal (MAB3580; Merck; 1:1,000); EPAC2 monoclonal (A-7, Santa Cruz, 1:200); CRHR1 polyclonal (EB08035; Everest Biotech; 1:500). Control and *Epac2*-shRNA constructs, expressing shRNA sequences and GFP, were previously described (Woolfrey *et al.*, 2009).

### Neuronal culture and transfections

Cortical neuronal cultures, derived from both sexes, were prepared from E18 Sprague-Dawley rat embryos (Srivastava *et al.*, 2011). Animals were habituated for 3 days before experimental procedures: experiments were carried out in accordance with the Home Office Animals (Scientific procedures) Act, United Kingdom, 1986. Cells were plated onto 18 mm glass coverslips (No 1.5), coated with poly-D-lysine (0.2 mg/mL, Sigma), at a density of 3×10^5^/well equal to 857/mm^2^. Neurons were cultured in feeding media: neurobasal medium (21103049) supplemented with 2% B27 (17504044), 0.5 mM glutamine (25030024) and 1% penicillin:streptomycin (15070063) (Life Technologies). Neuronal cultures were maintained in the presence of 200 µM D,L-amino-phosphonovalerate (APV, ab120004, Abcam) beginning on day 4 *in vitro* (DIV 4) in order to maintain neuronal health for long-term culturing (Srivastava *et al.*, 2011).

Primary cortical neurons were transfected with eGFP, control (scram-)RNAi or *Epac2*-RNAi at DIV 21, using Lipofectamine 2000 (Life Technologies). Briefly, 2-4 µg of plasmid DNA was mixed with Lipofectamine 2000 and incubated for 4 hours, before being replaced with fresh feeding media. Transfections were allowed to proceed for 2-5 days, after which cells were used for pharmacological treatment or immunocytochemistry.

### Pharmacological treatments of neuronal cultures

Treatments were performed in artificial cerebral spinal fluid (aCSF): (in mM) 125 NaCl, 2.5 KCL, 26.2 NaHCO_3,_ 1 NaH_2_PO_4_, 11 glucose, 5 HEPES, 2.5 CaCl_2,_ 1.25 MgCl_2_, and 0.2 APV. CRH was dissolved in H_2_O (10 mM), serially diluted to 1 µM in aCSF and applied directly to neuronal cultures at a final concentration of 100 nM. Final amount of H_2_O was < 0.01%; vehicle control was made up of H_2_O lacking compound. Treatments time was 30 minutes.

### Immunocytochemistry

Neurons were washed in phosphate-buffered saline (PBS), fixed in 4% formaldehyde/4% sucrose PBS for 10 minutes at room temperature, followed by incubation in methanol pre-chilled to −20°C for 10 minutes at 4°C. Fixed neurons were then permeabilized and blocked simultaneously (2% normal goat serum, 5425S, New England Biolabs, and 0.1% Triton X-100) before incubation in primary antibody solutions overnight and subsequent incubation with secondary antibodies the following day.

### Quantitative analysis of spine morphologies and immunofluorescence

Confocal images of double-stained neurons were acquired with a Leica SP-5 confocal microscope using a 63x oil-immersion objective (Leica, N.A. 1.4) as a z-series. Two-dimensional maximum projection reconstructions of images were generated and linear density calculated using ImageJ/Fiji (https://imagej.nih.gov/ij/) (Srivastava *et al.*, 2011). Morphometric analysis was performed on spines from two dendrites (secondary or tertiary branches), totaling 100 µm, from each neuron. The linear density and total gray value of each synaptic protein cluster were measured automatically using MetaMorph Software (Molecular Devices). Cultures directly compared were stained simultaneously and imaged with the same acquisition parameters. For each condition, 13-18 neurons from at least 3 separate experiments were used. Analyses were performed blind to condition and on sister cultures. In the green/magenta color scheme, co-localization is indicated by white overlap.

### Statistical analysis

All statistical analysis was performed in GraphPad. Differences in quantitative immunofluorescence and dendritic spine number were probed by one-way ANOVAs with Tukey correction for multiple comparisons. Error bars represent standard deviations in Figure 1D and standard errors of the mean in Figure 2B-C.

**Figure 1.**
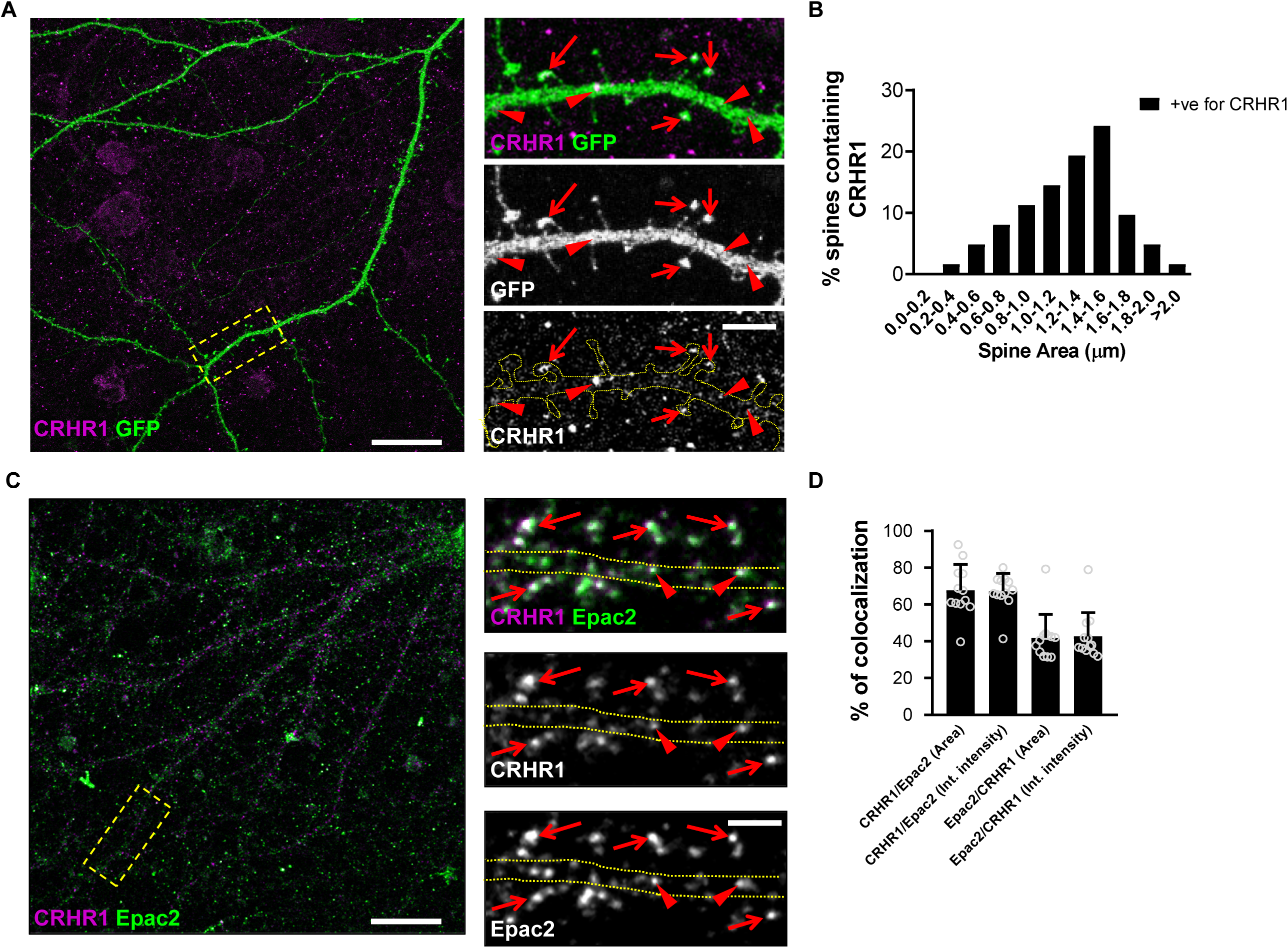
CRHR1 co-localizes with EPAC2 in primary cortical neurons. **(A)** Representative confocal microscopic images of a GFP-expressing cortical neuron double immunostained for GFP and CRHR1. The yellow box indicates the region of the dendrite displayed in magnified insets. Red arrows indicate spines enriched for CRHR1. Red arrowheads denote CRHR1 puncta within dendrites. **(B)** Histogram of the frequency of CRHR1 staining in spines of various sizes. The greatest enrichment of CRHR1 was observed in spines with an area of 1.0-1.6 µm. **(C)** Representative confocal microscopic images of a cortical neuron double immunostained for EPAC2 and CRHR1. The yellow box indicates the region of the dendrite displayed in magnified insets. Red arrows indicate spine-like structures where overlapping CRHR1 and EPAC2 puncta were observed. Red arrowheads denote co-localizing CRHR1 and EPAC2 puncta within dendrites. **(D)** Bar graph indicates quantitative measures of respective co-localization of immunofluorescent puncta. Scale bars: 20 µm for the main panels, 5 µm for the insets.

**Figure 2.**
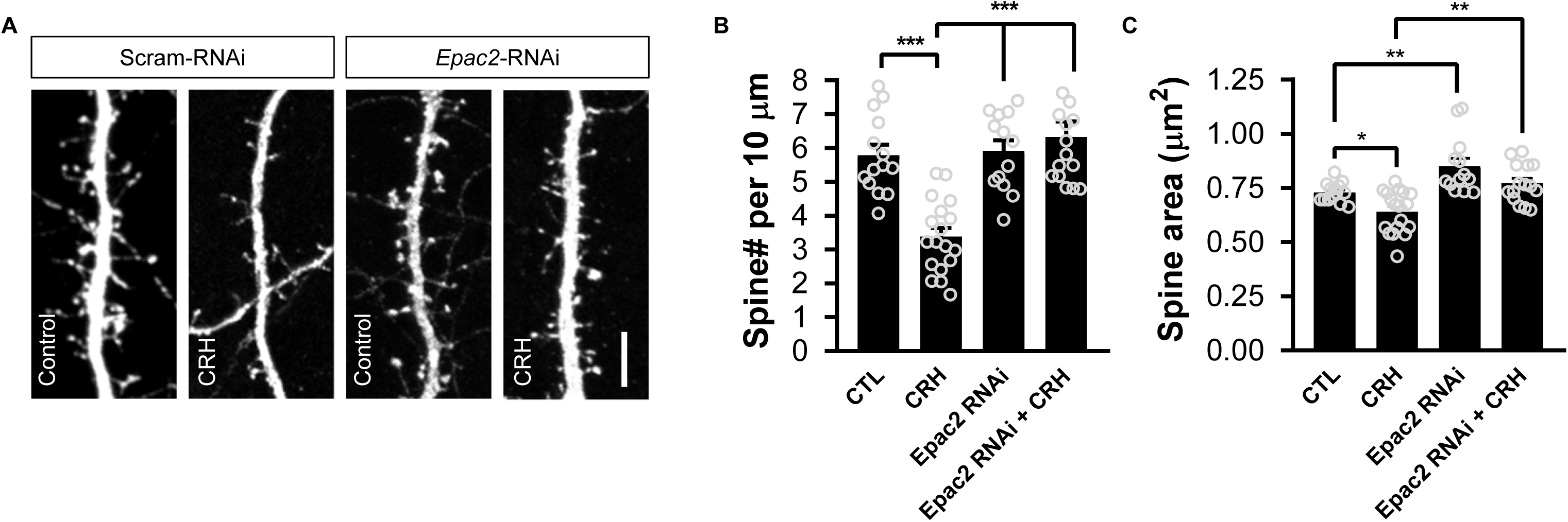
EPAC2 is required for CRH-induced rapid spine loss in primary cortical neurons. **(A)** Representative confocal microscopic images of dendrites from cortical neurons expressing either a control shRNA (Scram_RNAi) or *Epac2*-shRNA (Epac2_RNAi). Neurons were treated with either 100 nM CRH or vehicle control for 30 minutes. Scale bar: 5 µm. **(B)** Bar graph indicates quantitative measures of linear spine density. Treatment with CRH for 30 minutes induced a significant loss of spines (spines per 10 µm: control, 5.79±0.31; CRH, 3.39±0.25; *Epac2*-RNAi + control, 5.92±0.32; *Epac2*-RNAi + CRH, 6.33±0.45; F(3, 56) = 17.60, P<0.0001; Tukey post-hoc, ***=P<0.001). This effect was not observed in neurons in which EPAC2 was knocked down. **(C)** Quantitative measures of spine area reveals that CRH treatment results in a significant reduction of spine size; *Epac2-*shRNA increases spine size, whereas *Epac2-*shRNA expressing cells show no effect of CRH (spine area (µm^2^): control, 0.73±0.012; CRH, 0.64±0.023; *Epac2*-shRNA + control, 0.85±0.036; *Epac2*-shRNA + CRH, 0.77±0.023; F(3, 56) = 13.3, P<0.001; Tukey post-hoc, *=P<0.05, **=P<0.01).

## Results

### CRHR1 localizes to dendritic spines and co-localizes with EPAC2 in cortical neurons

Previous studies have shown that CRHR1 localizes in dendritic spines of CA1 hippocampal neurons (Chen *et al.*, 2010; Andres *et al.*, 2013). Whilst CRHR1 expression in cortical regions has been reported, whether or not this receptor is expressed in spines of cortical neurons has yet to be established. To investigate this, we immunostained DIV 25 primary cortical neurons expressing GFP for CRHR1 (**Figure 1A**). Immunoreactive puncta for CRHR1 could be observed within the somato-dendritic compartment of neurons. In individual neurons, CRHR1 clustered along dendrites, with prominent puncta evident near or at the base of dendritic spines (red arrowheads) and distinct clusters within spine heads (red arrows, **Figure 1A**). To determine whether CRHR1 localized to a specific subpopulation of dendritic spines, we classified spines containing CRHR1 according to dendritic spine area. Spines with an area of less than 1.0 µm were designated as “small”, whereas spines with an area of larger than 1.0 µm were designated as “large”. Of the spines that contained CRHR1, the majority (∼60%) were large (**Figure 1B**). These data indicate that in primary cortical neurons, CRHR1 is enriched at synapses of a subset of large dendritic spines.

Pharmacological studies have suggested that CRHR1 may signal via the EPAC proteins (Traver *et al.*, 2006; Grammatopoulos, 2012; Inda *et al.*, 2016). EPAC2 is the predominant EPAC protein expressed in cortical neurons with mature dendritic morphology, and is highly enriched in dendritic spines (Woolfrey *et al.*, 2009). Therefore, we reasoned that CRHR1 may colocalize with EPAC2 in cortical neurons. Endogenous EPAC2 and CRHR1 were detected by immunofluorescence using specific antibodies in DIV 23 primary cortical neurons. As previously described, EPAC2 was present along dendrites and in spines (**Figure 1C**) (Woolfrey *et al.*, 2009). Consistent with our data describing the localization of CRHR1 in cortical neurons, CRHR1 was also observed along dendrites and spine-like structures. Moreover, CRHR1 and EPAC2 were found to co-localize along dendrites (red arrowheads) and in a subset of spine-like structures (red arrows) (**Figure 1C**). Quantification of co-localization revealed that ∼70% of CRHR1 puncta co-localized with EPAC2, whereas only ∼40% of EPAC2 puncta colocalized with CRHR1 (**Figure 1D**). Taken together, these data indicate that CRHR1 is ideally positioned to interact with and signal via EPAC2 at synapses.

### CRH-mediated spine loss is dependent on EPAC2 in cortical neurons

As our data indicated that CRHR1 co-localizes with EPAC2, we hypothesized that CRH-mediated spine loss may be mediated by this Rap GEF. To test this, we treated primary cortical neurons expressing a control shRNA (scram-RNAi)or an shRNA specific for *Epac2* (*Epac2*-RNAi; **Supplemental Figure 1**) (Woolfrey *et al.*, 2009) with CRH. Consistent with previous reports, 30 minutes of CRH exposure resulted in a significant loss of dendritic spines (**Figure 2**). *Epac2*-RNAi alone had no effect on spine linear density;.however, treatment of cortical neurons expressing *Epac2*-RNAi with CRH no longer resulted in a reduction in dendritic spine density (**Figure 2B**). Examination of spine morphology revealed that CRH treatment resulted in an overall decrease in spine size; this effect was no longer evident in neurons treated with CRH and expressing *Epac2*-RNAi (**Figure 2C**). Taken together these data suggest that CRH signals via EPAC2 to induce the rapid loss of dendritic spines.

## Discussion

Previous studies have shown that CRH signaling via CRHR1 can cause the rapid and persistent loss of dendritic spines in hippocampal neurons (Chen *et al.*, 2008; Chen *et al.*, 2010). This loss of spine density is further correlated with memory defects associated with acute stress (Chen *et al.*, 2010). In this study, we build upon these findings to show that in primary cortical neurons, CRHR1 localizes to synapses in dendritic spines, where it co-localizes with the Rap GEF EPAC2. Moreover, acute exposure to CRH resulted in the rapid loss of dendritic spines, an effect that was attenuate by the knockdown of *Epac2*. Taken together, these data indicate that CRH signaling via a CRHR1/EPAC2-dependent signaling pathway is responsible for the actions of this hormone in regulating the density of dendritic spines and contributing to acute stress effects.

Stress is a biologically important event that can have both positive and negative effects on brain function. Multiple lines of evidence have demonstrated that stress can induce a range of morphological changes in neuronal and synaptic structure. CRH is released in response to stress, and recent findings have shown that blocking the CRHR1 receptor was sufficient to block stress-induced spine loss (Chen *et al.*, 2008; Chen *et al.*, 2010). Interestingly, in hippocampal neurons, CRH-mediated spine loss on CA1 hippocampal neurons is dependent on synaptic-activity (Andres *et al.*, 2013). Data indicates that EPAC2 is required for CRH-dependent spine loss on cortical neurons. Whether CRH acts via the regulation of both EPAC2 and synaptic-activity to induce spine loss, or whether these are independent mechanisms dependent on cell type is currently unclear.

Moreover, recent studies have shown that EPAC2 expression is increased in response to acute stress, and is involved in controlling cellular responses to acute stress (Aesoy *et al.*, 2018). Indeed, animals lacking the EPAC proteins display increased anxiety and depressive behaviors (Srivastava *et al.*, 2012; Yang *et al.*, 2012). Interestingly, our previous work has shown that EPAC2 is a key regulator of dendritic spine structural plasticity in response to a number of extrinsic stimuli (Woolfrey *et al.*, 2009; Srivastava *et al.*, 2012). Consistent with these studies, our current data indicate that EPAC2 is required for mediating CRH-induced spine loss. In hippocampal neurons, CRH-mediated spine loss is dependent on RhoA activity (Chen *et al.*, 2013) as well as a nectin-3/afadin complex (Wang *et al.*, 2013). Afadin is a direct target of Epac-Rap1 signaling and, a Rap1-afadin complex has been shown to regulate RhoA activity and cytoskeletal dynamics in endothelia cells (Birukova *et al.*, 2013). Thus, an intriguing possibility is that these signaling molecules cooperate to induced spine loss following CRH-activity. Interestingly, it has recently been shown that CRHR1 engages atypical soluble adenylate cyclase to signal to EPAC proteins (Inda *et al.*, 2016) indicating an indirect interaction between these 2 proteins.

It is also interesting to note that EPAC2 has been associated with anxious and depressive behaviors as well as in stress responses (Yang *et al.*, 2012; Zhou *et al.*, 2016; Aesoy *et al.*, 2018). Furthermore, EPAC2 plays an important role in cognitive function, including learning and memory (Srivastava *et al.*, 2012; Yang *et al.*, 2012). Given that CRH and CRHR1 have strongly been implicated in stress-mediated effects on these cognitive functions (Maras & Baram, 2012; Henckens *et al.*, 2016), the data presented in this study suggest that a CRH/CRHR1/EPAC2 pathway may be critical for these effects. Taken together, these data reveal a novel mechanism involving EPAC2, by which CRH-induced rapid modulation of dendritic spines can occur. Future studies will be required to understand whether and how this pathway is involved in mediating responses to chronic stress at both the morphological and behavioral levels.

## Acknowledgments

We thank Tallie Baram for her helpful discussion on this study. This work was supported by grants from the Medical Research Council, MR/L021064/1, Royal Society UK (Grant RG130856), and the Brain and Behavior Foundation (formally National Alliance for Research on Schizophrenia and Depression (NARSAD); Grant No. 25957), awarded to D.P.S.; and NIH grants R01MH071316 and R01MH097216 to P.P.

## Author Contributions

P.P and D.P.S. were responsible for the conception and design of the work. Z.X. and D.P.S. were responsible for data collection, analysis, and interpretation.

Drafting and critical revision of the article was carried out by P.P. and D.P.S.

Final approval of the version to be published was confirmed by Z.X., P.P. and D.P.S.

## Data Accessibility Statement

All primary data are available upon request from the authors.

## Abbreviations

aCSF: artificial cerebral spinal fluid;
APV: amino-phosphonovalerate;
CRH: corticotropin-releasing hormone;
CRHR1: corticotropin-releasing hormone receptor 1;
DIV: days *in vitro*;
GEF: guanine nucleotide exchange factor;
GFP: green fluorescent protein;
PBS: phosphate-buffered saline

## References

Aesoy, R., Muwonge, H., Asrud, K.S., Sabir, M., Witsoe, S.L., Bjornstad, R., Kopperud, R.K., Hoivik, E.A., Doskeland, S.O. & Bakke, M. (2018) Deletion of exchange proteins directly activated by cAMP (Epac) causes defects in hippocampal signaling in female mice. PloS one, 13, e0200935.

Andres, A.L., Regev, L., Phi, L., Seese, R.R., Chen, Y., Gall, C.M. & Baram, T.Z. (2013) NMDA receptor activation and calpain contribute to disruption of dendritic spines by the stress neuropeptide CRH. The Journal of neuroscience : the official journal of the Society for Neuroscience, 33, 16945–16960.

Birukova, A.A., Tian, X., Tian, Y., Higginbotham, K. & Birukov, K.G. (2013) Rap-afadin axis in control of Rho signaling and endothelial barrier recovery. Mol Biol Cell, 24, 2678–2688.

Chen, Y., Dube, C.M., Rice, C.J. & Baram, T.Z. (2008) Rapid loss of dendritic spines after stress involves derangement of spine dynamics by corticotropin-releasing hormone. The Journal of neuroscience : the official journal of the Society for Neuroscience, 28, 2903–2911.

Chen, Y., Kramar, E.A., Chen, L.Y., Babayan, A.H., Andres, A.L., Gall, C.M., Lynch, G. & Baram, T.Z. (2013) Impairment of synaptic plasticity by the stress mediator CRH involves selective destruction of thin dendritic spines via RhoA signaling. Mol Psychiatry, 18, 485–496.

Chen, Y., Rex, C.S., Rice, C.J., Dube, C.M., Gall, C.M., Lynch, G. & Baram, T.Z. (2010) Correlated memory defects and hippocampal dendritic spine loss after acute stress involve corticotropin-releasing hormone signaling. Proceedings of the National Academy of Sciences of the United States of America, 107, 13123–13128.

Grammatopoulos, D.K. (2012) Insights into mechanisms of corticotropin-releasing hormone receptor signal transduction. British journal of pharmacology, 166, 85–97.

Henckens, M.J., Deussing, J.M. & Chen, A. (2016) Region-specific roles of the corticotropin-releasing factor-urocortin system in stress. Nature reviews. Neuroscience, 17, 636–651.

Inda, C., Dos Santos Claro, P.A., Bonfiglio, J.J., Senin, S.A., Maccarrone, G., Turck, C.W. & Silberstein, S. (2016) Different cAMP sources are critically involved in G protein-coupled receptor CRHR1 signaling. J Cell Biol, 214, 181–195.

Maras, P.M. & Baram, T.Z. (2012) Sculpting the hippocampus from within: stress, spines, and CRH. Trends in neurosciences, 35, 315–324.

Sanders, J. & Nemeroff, C. (2016) The CRF System as a Therapeutic Target for Neuropsychiatric Disorders. Trends in pharmacological sciences, 37, 1045–1054.

Srivastava, D.P., Jones, K.A., Woolfrey, K.M., Burgdorf, J., Russell, T.A., Kalmbach, A., Lee, H., Yang, C., Bradberry, M.M., Wokosin, D., Moskal, J.R., Casanova, M.F., Waters, J. & Penzes, P. (2012) Social, communication, and cortical structural impairments in Epac2-deficient mice. The Journal of neuroscience : the official journal of the Society for Neuroscience, 32, 11864–11878.

Srivastava, D.P., Woolfrey, K.M. & Penzes, P. (2011) Analysis of dendritic spine morphology in cultured CNS neurons. Journal of visualized experiments : JoVE, e2794.

Traver, S., Marien, M., Martin, E.,Hirsch, E.C. & Michel, P.P. (2006) The phenotypic differentiation of locus ceruleus noradrenergic neurons mediated by brain-derived neurotrophic factor is enhanced by corticotropin releasing factor through the activation of a cAMP-dependent signaling pathway. Molecular pharmacology, 70, 30–40.

Wang, X.D., Su, Y.A., Wagner, K.V., Avrabos, C., Scharf, S.H., Hartmann, J., Wolf, M., Liebl, C., Kuhne, C., Wurst, W., Holsboer, F., Eder, M., Deussing, J.M., Muller, M.B. & Schmidt, M.V. (2013) Nectin-3 links CRHR1 signaling to stress-induced memory deficits and spine loss. Nature neuroscience, 16, 706–713.

Woolfrey, K.M., Srivastava, D.P., Photowala, H., Yamashita, M., Barbolina, M.V., Cahill, M.E., Xie, Z., Jones, K.A., Quilliam, L.A., Prakriya, M. & Penzes, P. (2009) Epac2 induces synapse remodeling and depression and its disease-associated forms alter spines. Nature neuroscience, 12, 1275–1284.

Yang, Y., Shu, X., Liu, D., Shang, Y., Wu, Y., Pei, L., Xu, X., Tian, Q., Zhang, J., Qian, K., Wang, Y.X., Petralia, R.S., Tu, W., Zhu, L.Q., Wang, J.Z. & Lu, Y. (2012) EPAC null mutation impairs learning and social interactions via aberrant regulation of miR-124 and Zif268 translation. Neuron, 73, 774–788.

Zhou, L., Ma, S.L., Yeung, P.K., Wong, Y.H., Tsim, K.W., So, K.F., Lam, L.C. & Chung, S.K. (2016) Anxiety and depression with neurogenesis defects in exchange protein directly activated by cAMP 2-deficient mice are ameliorated by a selective serotonin reuptake inhibitor, Prozac. Translational psychiatry, 6, e881.

