## Supplementary figures and images for "EPAC2 is required for corticotropin-releasing hormone-mediated spine loss"

### Supplemental Figure 1

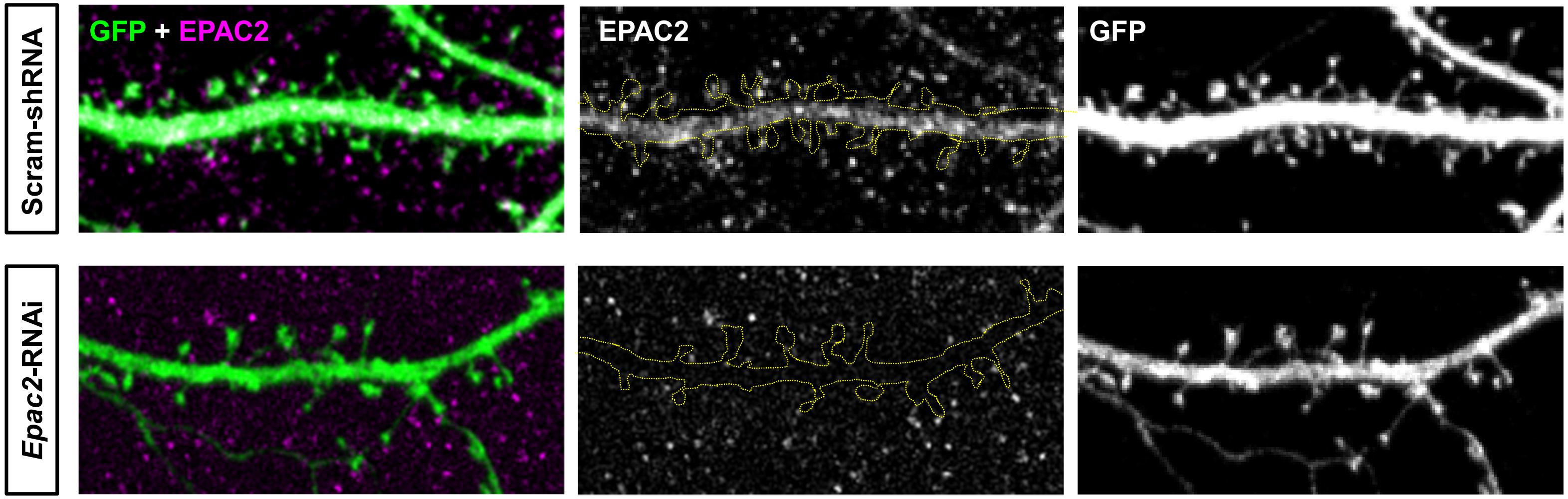
